# The Indirect Pathway of the Basal Ganglia Promotes Negative Reinforcement, But Not Motor Suppression

**DOI:** 10.1101/2022.05.18.492478

**Authors:** Brian R. Isett, Katrina P. Nguyen, Jenna C. Schwenk, Christen N. Snyder, Kendra A. Adegbesan, Ugne Ziausyte, Aryn H. Gittis

## Abstract

Optogenetic stimulation of Adora2a receptor expressing spiny projection neurons (A2A-SPN) in the striatum drives locomotor suppression and negative reinforcement, results attributed to activation of the indirect pathway. The sole long-range projection target of A2A-SPNs is the external globus pallidus (GPe). Unexpectedly, we found that inhibition of the GPe did not suppress movement, but did drive robust negative reinforcement in a real-time place preference assay. Within the striatum, A2A-SPNs inhibit other SPNs through a short-range inhibitory collateral network, and we found that optogenetic stimuli that drove motor suppression shared a common mechanism of recruiting this inhibitory collateral network. Our results suggest that the indirect pathway plays a more prominent role in negative reinforcement than in motor control and challenges the assumption that activity of A2A-SPNs is synonymous with indirect pathway activity.

## Introduction

The direct and indirect pathways of the basal ganglia are modeled as separate, parallel pathways that converge at the level of basal ganglia output (Albin et al., 1989; DeLong, 1990). Activity along the direct pathway is associated with increased locomotion and positive reinforcement, whereas activity along the indirect pathway is associated with reduced locomotion and negative reinforcement/punishment (Cox and Witten, 2019; Gerfen and Surmeier, 2011; Kravitz and Kreitzer, 2012; Nelson and Kreitzer, 2014). Views about the behavioral effects of the direct and indirect pathways were first developed through anatomical and clinical observations, and later refined through the use of optogenetics, which enabled targeting of molecularly defined neuronal subpopulations in the striatum (Gerfen et al., 1990; Kravitz et al., 2010; Lenz and Lobo, 2013; Parker et al., 2016; Soares-Cunha et al., 2016).

The indirect pathway is typically targeted using mice where Cre recombinase is expressed under control of the Adora2a receptor (A2A-Cre) (Gong et al., 2007; Schiffmann et al., 1991). Axons of A2A-SPNs inhibit neurons in the external globus pallidus (GPe), and through a series of subsequent connections leads to disinhibition of basal ganglia output nuclei, giving rise to the indirect pathway (Cazorla et al., 2014; Freeze et al., 2013; Kovaleski et al., 2020; Pollack et al., 1993; Sano et al., 2013; Smith et al., 1998). Optogenetic stimulation of A2A-SPNs decreases movement (Bakhurin et al., 2020; Kravitz et al., 2010; Roseberry et al., 2016), a result widely interpreted as evidence of the indirect pathway’s role in motor suppression. Optogenetic stimulation of A2A-SPNs has also been shown to drive negative reinforcement, at thresholds even lower than those needed to inhibit locomotion (Kravitz et al., 2012; LeBlanc et al., 2020; Yttri and Dudman, 2016). These multifaceted behavioral effects of A2A-SPN stimulation have been attributed to engagement of distinct motor and limbic loops of the indirect pathway (Alexander et al., 1986; Lee et al., 2020; Lynd‐Balta and Haber, 1994).

The GPe is the sole target of extra-striatal projections of A2A-SPNs, and GPe neurons encode information related to both movement and reward (Arkadir et al., 2004; Kim et al., 2017; Schechtman et al., 2016). Although the GPe is an obligate node for striatal signals to travel along the indirect pathway, pharmacological inhibition or lesions of the GPe have been found to produce only modest effects on movement, ranging from a 10-15% decrease in locomotion (Lilascharoen et al., 2021) to increased muscle flexion (Kato and Kimura, 1992), to no effect at all (Horak and Anderson, 1984; Soares et al., 2004). And although the GPe encodes information about reward and value, it is unclear whether these signals are sufficient to drive behavior.

Optogenetic stimulation of A2A-SPNs inhibits GPe neurons (Cazorla et al., 2014), but has also been shown to inhibit other SPNs through inhibitory collaterals in the striatum (Dobbs et al., 2016; Kravitz et al., 2013; Taverna et al., 2008; Tepper et al., 2008). The simultaneous activation of these two circuits creates ambiguity for experimental interpretation, and merits a closer assessment of which behaviors are driven through the canonical indirect pathway (downstream inhibition of the GPe), and which behaviors are generated through collateral inhibition within the striatum.

To address this question, we used optogenetics, combined with *in vivo* recordings, to establish the circuits responsible for the locomotor vs. reinforcing effects of A2A-SPN stimulation. We find that both direct and synaptic inhibition of the GPe drives negative reinforcement, but does not inhibit movement. Suppression of movement was only observed under stimulus conditions in which the inhibitory collaterals in the striatum were also recruited during stimulation. Our results suggest that negative reinforcement, not motor suppression, is the primary behavioral correlate of signaling along the indirect pathway, and that the motor suppressing effects of A2A-SPN stimulation are driven through inhibitory collaterals within the striatum.

## Results

### A2A-SPN Stimulation Inhibits Movement and Drives Real Time Place Avoidance

Optogenetic stimulation of A2A-SPNs in the striatum is associated with effects on both locomotion and reinforcement. To quantify these behaviors, A2A-Cre mice expressing channelrhodopsin (ChR2) were placed in a 50 cm^2^ open field arena and optogenetic stimulation was either delivered in an open loop or closed loop configuration to study its effects on movement or reinforcement, respectively. In open loop experiments, bilateral stimulation was delivered for 10 sec (1 mW, continuous light) every 3 min. for a total of 10 trials. Optogenetic stimulation of A2A-SPNs robustly decreased movement, quantified as a decrease in average speed (paired t-test, p<0.001, N=15 mice) (Fig. 1A-C). This effect was driven primarily by a reduction in the number of movement bouts (p<0.0001, N=15 mice) (Fig. 1E). The duration of movement bouts and speed within a bout were not affected by stimulation (Fig. 1E). No movement effects were observed in control A2A-Cre mice expressing EYFP instead of ChR2 (Fig. 1D).

**Figure 1:**
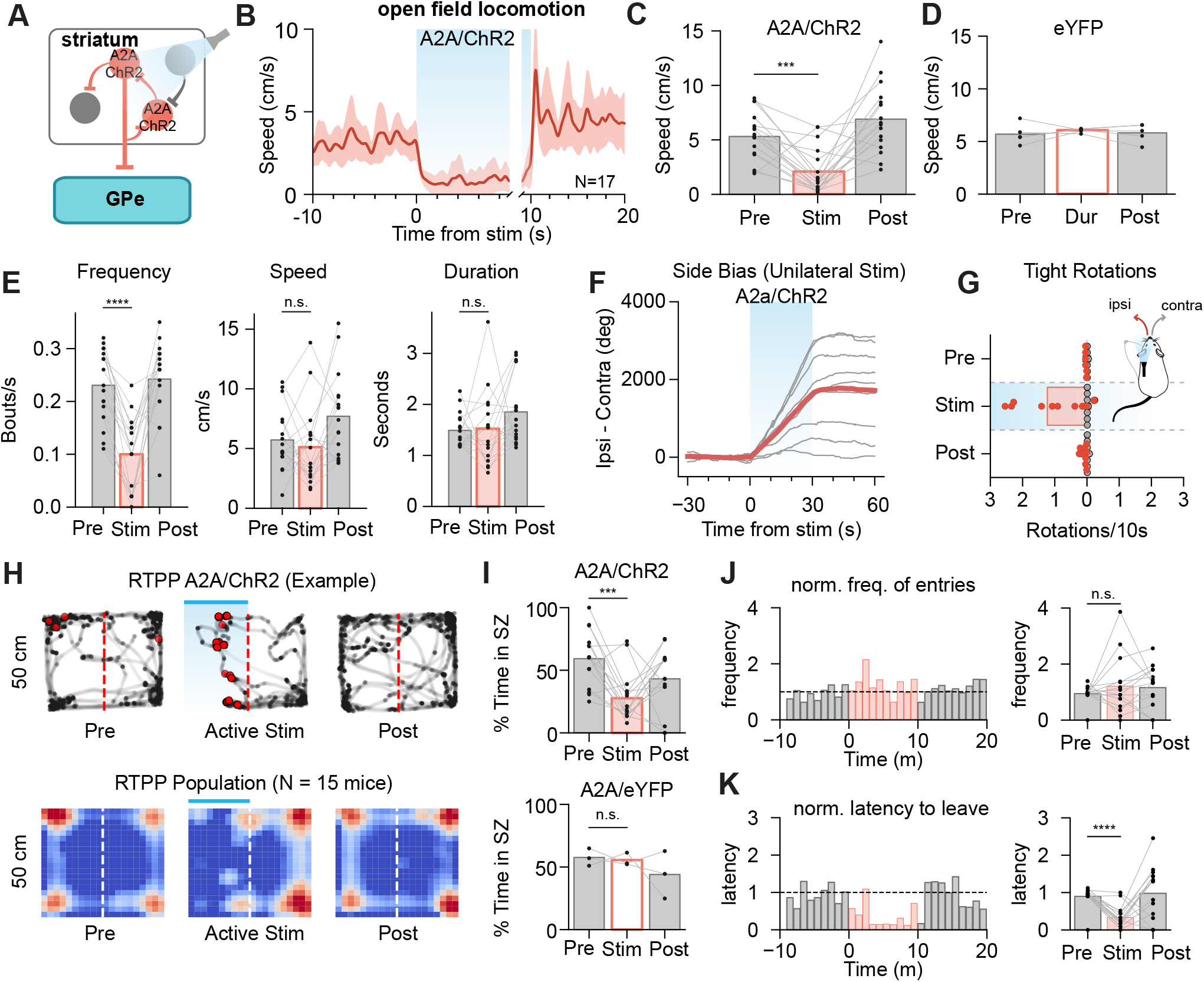
A2A-SPN excitation reduces locomotion, causes avoidance, and induces ipsilateral rotations. **A**. Circuit schematic of experiment. **B**. Instantaneous locomotor speed in the open field, averaged across N = 17 mice. Median (dark line) ± sem (shaded area). To average across mice who received either 10 s or 30 s of stimulation, movement traces were truncated after the first 10 s of stimulation, and then aligned to the offset of stimulation (x-axis break). **C**. A2A-SPN stimulation drove a significant reduction in speed (pre: 5.31 ± 0.51 cm/s vs stim: 2.07 ± 0.49 cm/s; paired t-test: p<0.001, N=17). Data from individual mice, and population averages are shown. **D**. eYFP controls showed no significant changes in speed (pre: 5.69 ± 0.38 cm/s vs stim: 6.04 ± 0.08 cm/s; paired t-test: p=0.48). **E**. Stimulation decreased the frequency of ambulation bouts (pre: 0.23 ± 0.02 bouts/s vs stim: 0.10 ± 0.02 bouts/s; paired t-test: p<0.0001), but not ambulation bout speed (pre: 5.73 ± 0.61 cm/s vs stim: 5.12 ± 0.79 cm/s; paired t-test: p=0.50, N=17) or duration (pre: 1.50 ± 0.08 sec vs stim: 1.52 ± 0.18 sec; paired t-test: p=0.93, N=17). **F**. Accumulated angle displacement before, during, and after A2A-SPN stimulation for individual mice (thin lines) and the population average (thick line). Time of stimulation is indicated with shaded blue bar. **G**. A2A-SPN stimulation induced tight rotations towards the side ipsilateral to stimulation (*see methods*). **H**. *top:* Movement path from a representative mouse during the RTPP task. Red circles denote periods of immobility. *bottom*: Heat maps averaged across all mice (N = 17) displaying the % time spent in each location of the open field (20×20 grid). **I**. *top:* Mice avoided the side of stimulation in a real time place preference (RTPP) task (pre: 59.2 ± 5.55% vs stim: 27.7 ± 4.97%; paired t-test: p=0.0007, N=15). *bottom:* Mice injected with eYFP (N=3) showed no place preference (pre: 57.63 ± 3.28% vs stim: 55.54 ± 2.43%; paired t-test, p=0.78). **J**. The frequency of stimulation zone entrances in 1-minute bins, normalized to the average baseline period value for each mouse. Colored bars represent the active stimulation period, in which crossing over to the stimulated zone triggered bilateral optogenetic stimulation. The frequency of stimulation zone entries did not change (0.94 ± 0.07 pre versus 1.21 ± 0.26 during; p=0.32; N=15). **K)** The latency to leave the stimulated zone in 1-minute bins across the trial, normalized to the average baseline period value for each mouse. Latency to exit the stimulation zone significantly decreased during stimulation (0.90 ± 0.06 pre versus 0.34 ± 0.07 during; p<0.0001; N=15). Mean ± sem unless otherwise noted.

Unilateral stimulation of A2A-SPNs biased movement towards the side of stimulation (Fig. 1F), and in some cases drove tight, full-body ipsilateral rotations (Fig. 1G). As shown in Fig. 1F, before stimulation, mice did not have a turning bias, but during stimulation, mice frequently turned or walked towards the side of stimulation, leading to accumulated angle displacement. At the offset of stimulation, this bias corrected, and mice no longer showed a side bias.

To study the effects of A2A-SPN stimulation on reinforcement, we used a real-time place preference assay (RTPP) (Kravitz and Kreitzer, 2012; Kravitz et al., 2012), where closed loop stimulation (1 mW, continuous light) was triggered when mice entered one half of the arena (stimulation zone). Mice were run on two different days with the stimulation zone switching from one side to the other across days to avoid intrinsic side biases. During baseline exploration, before stimulation, mice spent 59.2 ± 5.55% of their time in the inactive stimulation zone, but during the active stimulation period, that time dropped to 27.7± 4.97 % (p=0.0007, N=15). Avoidance of the stimulated side quickly abated after the task, consistent with the lack of long carry-over effects following A2A-SPN stimulation (Kravitz et al., 2012; Yttri and Dudman, 2016). During a 10 minute post-stimulation washout period, mice were allowed to freely explore the arena with no further stimulation, and they quickly resumed exploration of both halves of the arena (Fig. 1I, *top*). Mice expressing a control eYFP fluorophore, rather than ChR2, showed no place preference in response to closed-loop light stimulation (Fig. 1I, *bottom*).

To determine whether the real-time place avoidance produced by A2A-SPN stimulation was due to fewer entries into the stimulated side *(“frequency of entries”*, Fig. 1J), or less time spent in the stimulation zone once entered *(“latency to leave”*, Fig. 1K), we considered each of these variables independently. To control for potential movement effects of A2A-SPN stimulation during the active stimulation period, latency to leave was calculated as the total time mice spent mobile in the stimulation zone before leaving. As shown, the normalized frequency that mice entered the stimulated zone did not change (0.94 ± 0.07 pre versus 1.21 ± 0.26 during; p=0.32; N=15), but they were much quicker to leave the zone during periods of active stimulation (0.90 °10.06 pre versus 0.34 ± 0.07 during; p<0.0001; N=15). Taken together, these results demonstrate that A2A-SPN stimulation induces real-time place avoidance by driving mice to leave the zone quickly upon entering, rather than learning to avoid entering the zone.

### GPe Inhibition Drives Real Time Place Avoidance But Not Motor Suppression

The behavioral effects produced by A2A-SPN stimulation, as described above, are attributed to effects of the indirect pathway. The first synaptic connection of the indirect pathway is an inhibitory projection from A2A-SPNs to the GPe, and the GPe is the sole recipient of A2A-SPN projections outside of the striatum. To measure the responses of GPe neurons to optogenetic stimulation of A2A-SPNs, we performed *in vivo* recordings from awake, head fixed mice (Fig. 2A-B). During optogenetic stimulation of A2A-SPNs (1 mW, 10 s continuous light) 68.5% of GPe neurons were inhibited (63/92 units across 3 mice) (Fig. 2D). On average, z-scored firing rates were decreased by -8.08 ± 3.44 standard deviations of baseline firing rates during stimulation (Fig. 2C). Of the remaining 29 GPe units that were not inhibited by A2A-SPN stimulation, 27.2% showed no significant modulation, and 1.1% were excited (Fig. 2D).

**Figure 2:**
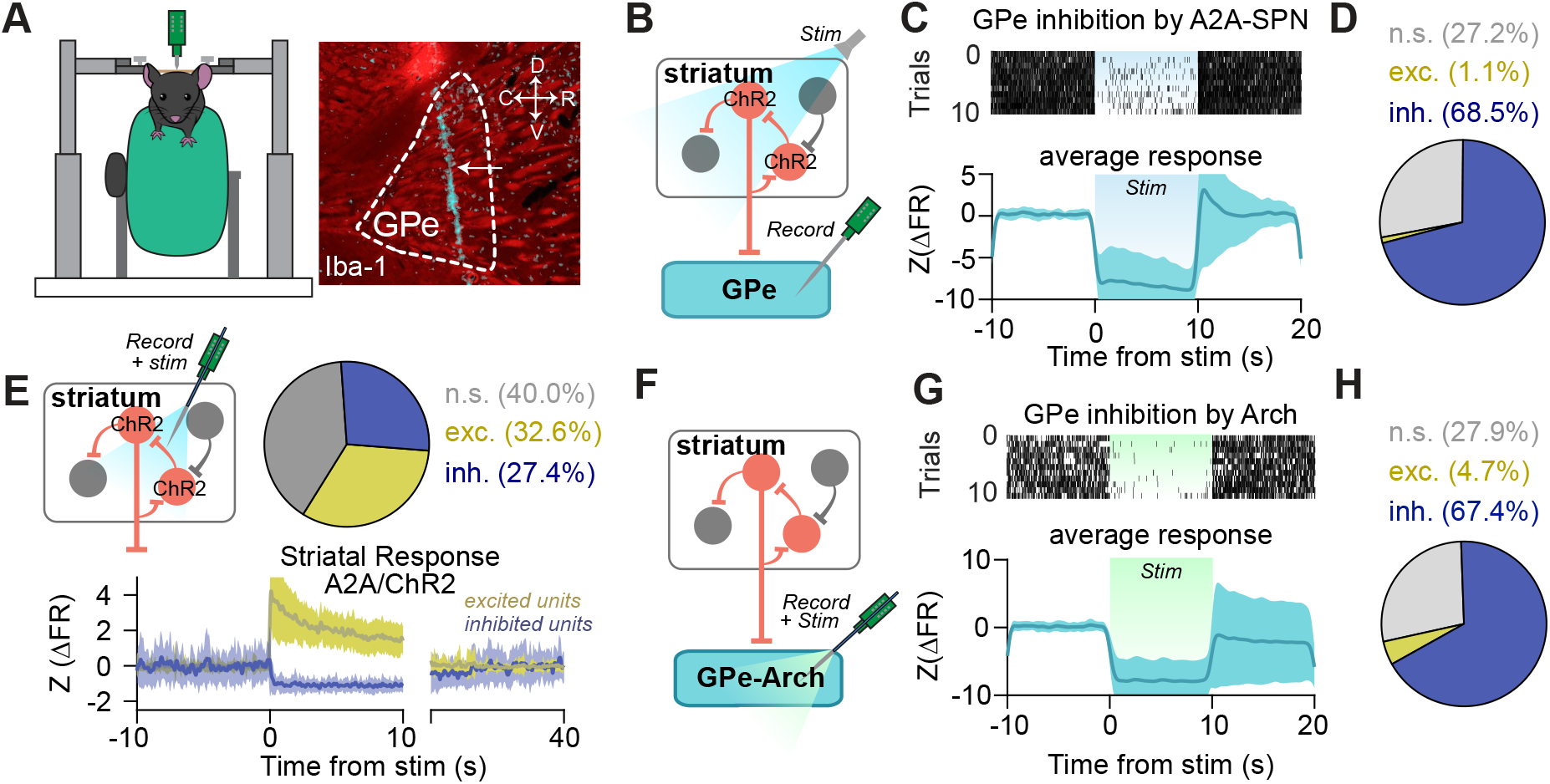
Physiological effects of A2A-SPN stimulation in the GPe and striatum. **A**. *In vivo* head-fixed running wheel setup. Example histology verifying the location of the recording tract in the GPe. **B**. A2A-cre mice received light stimulation in the striatum while units were recorded in the GPe. **C**. *top:* sample GPe unit inhibited by A2A-SPN soma stimulation in the striatum. *bottom:* Average population response to A2A-SPN stimulation represented as z-scored PSTH. Data aligned to the onset of stimulation. **D**. Summary of GPe response to A2A-SPN stimulation: 27.2% n.s. (n=25/92), 1.1% excited (n=1/92), 68.5% inhibited (n=63/92). **E**. Simultaneous stimulation of A2A-SPN and recording in the striatum with an optrode showed both excitatory (39%, n=55/143) and inhibitory (14.6%, n=21/143) responses. **F**. Direct inhibition of the Gpe with CAG-Arch. **G**. *top:* sample Gpe unit inhibited with an optrode using green light at 1mW for 10 s. *bottom:* Average population response to green light stimulation of Arch-expressing Gpe neurons represented as z-scored PSTH. **H)** Summary of GPe response when directly inhibited using CAG-Arch: 27.9% n.s. (n=12/43), 4.7% excited (n=2/43), 67.4% inhibited (n=29/43). Throughout the figure, shaded area denotes sem.

Although the GPe is the only extra-striatal target of A2A-SPNs, SPNs are known to laterally inhibit each other (Burke et al., 2017; Dobbs et al., 2016; Taverna et al., 2008) (Fig. 2E). To quantify the strength of this collateral inhibition, we recorded neural activity in the striatum during optical stimulation of A2A-SPNs using an optrode. Optogenetic stimulation within the striatum resulted in excitation of 31/95 units (32.6%, across 3 mice), and inhibition of 26/95 units (27.4%, across 3 mice) (Fig. 2E). On average, the z-score of inhibited neurons decreased by by -1.07 ± 0.45 standard deviations. Taken together, these results confirm that a conventional optogenetic protocol used to stimulate A2A-SPNs drives simultaneous inhibition in the striatum, as well as the GPe. This raises the question of which neuronal responses are responsible for the behaviors seen during A2A-SPN stimulation?

To disambiguate the behavioral effects of these two targets we bypassed the striatum and inhibited the GPe directly using archaerhodopsin (CAG-Arch) (Fig. 2F). This approach was validated using an optrode to record the responses of Arch-expressing GPe neurons during green light stimulation (1 mW, 10 s continuous). Optogenetic inhibition resulted in 67% of GPe neurons showing significant inhibition (29/43 across 3 mice) (Fig. 2H), where z-scored firing rates dropped to -7.60 ± 3.22 standard deviations of baseline firing rates (Fig. 2G). Of the remaining 14 GPe units that were not inhibited by green light, 27.9% showed no significant modulation, and 4.7% were excited (Fig. 2H). The magnitude of firing rate inhibition and fraction of neurons affected was similar to that observed following optogenetic stimulation of A2A-SPNs (A2A-SPN: 8.08 ± 3.44 std. dev. vs GPe: 7.60 ± 3.22 std. dev.; Wilcoxon rank sum test; p=0.38).

Next, we repeated the same battery of behavioral tasks used to evaluate the effects of A2A-SPN stimulation, but this time inhibited the GPe directly (Fig. 3A). Contrary to predicted effects of indirect pathway stimulation, inhibition of the GPe (1 mW continuous light for 10 sec, 10 trials spaced 3 min. apart) did not decrease speed, but rather produced a slight increase in speed (Fig. 3B-C). This effect was driven primarily by an increase in locomotor speed during movement bouts and an increase in bout duration (Fig. 3E). No change was seen in the frequency of bout initiation (Fig. 3E).

**Figure 3.**
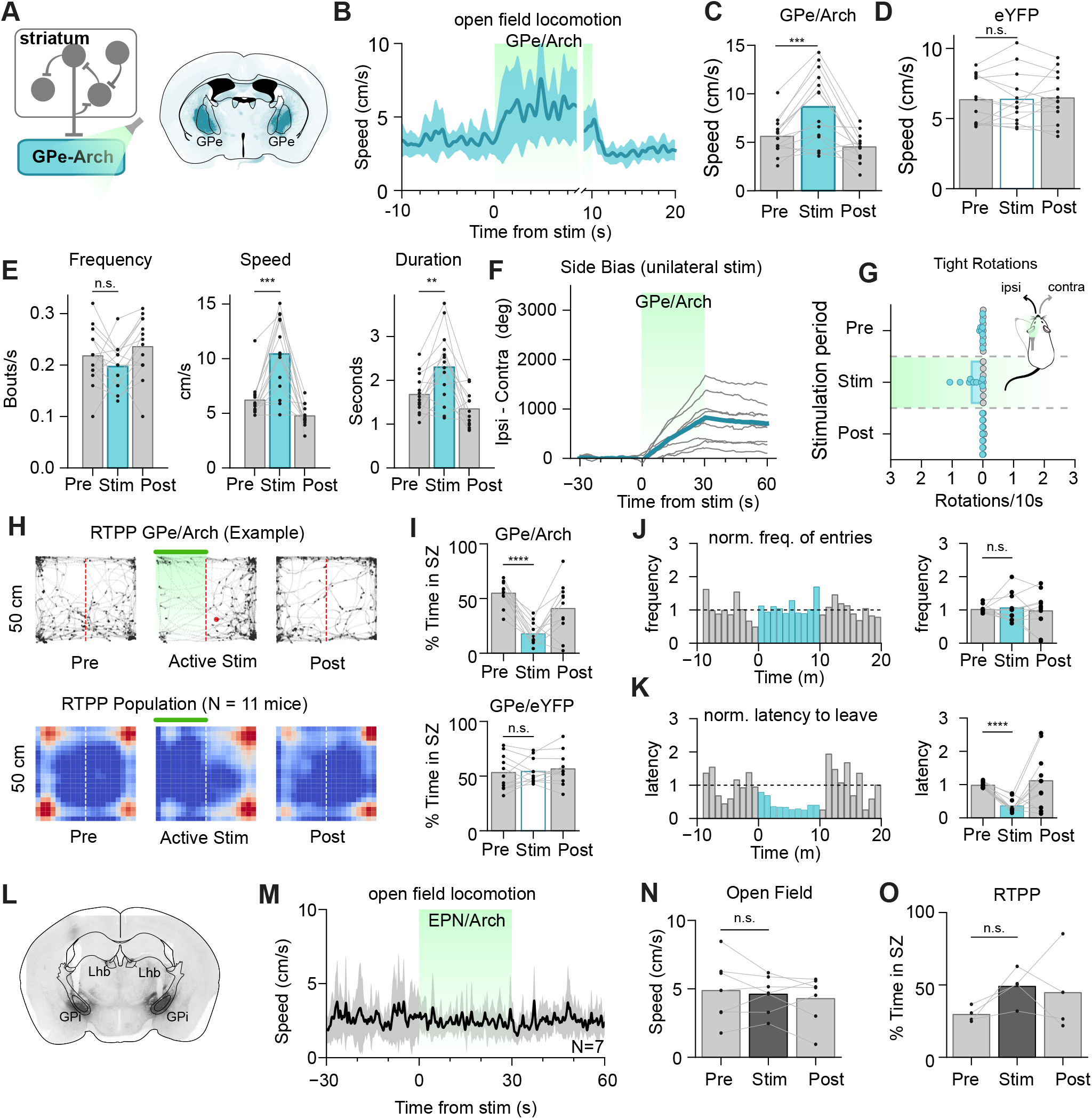
Inhibiting GPe activity does not reduce locomotion, but does cause avoidance. **A**. Circuit schematic of behavioral manipulation. **B**. Average (N=14) instantaneous speed in the open field, aligned to onset of stimulation. Median (dark line) ± sem (shaded area). **C**. Optogenetic inhibition of GPe increased speed in an open field (pre: 5.51±0.47 cm/s vs stim: 8.50±0.95 cm/s; paired t-test, p=0.0004, N=14). **D**. Inhibition of GPe in eYFP mice showed no effects of stimulation on speed (pre: 6.34±0.45 cm/s vs stim: 6.37±0.51 cm/s; paired t-test, p=0.93, N=12). **E**. Bilateral optogenetic inhibition of all GPe neurons significantly increased ambulation bout speed (pre: 6.20±0.44 cm/s vs stim: 10.43±0.88 cm/s; paired t-test, p=0.0001) and duration (pre: 1.68±0.11 sec vs stim: 2.30±0.21 sec; paired t-test, p=0.003), but not bout rate (pre: 0.22 ± 0.01 bouts/s vs stim: 0.20 ± 0.01 bouts/s; paired t-test, p=0.21). **F**. Accumulated angle displacement before, during, and after GPe inhibition for individual mice (thin lines) and the population average (thick line). **G**. Inhibiting the GPe did not drive strong ipsilateral rotations. **H**. Representative *(top)* and population average *(bottom)* data on the RTPP assay where entry into the stimulated zone trigged optogenetic inhibition of the GPe. Red points represent instances of freezing, or a movement slow-down. **I**. *top:* Mice expressing Arch in the GPe avoided the side of stimulation (pre: 54.8 5± 3.22% vs stim: 17.72 ± 2.91%; paired t-test, p<0.0001, N=11*). bottom:* mice expressing eYFP showed no place preference to the stimulated zone (pre: 52.2 ± 4.6% vs stim: 54.2 ± 3.2%; paired t-test, p=0.80, N=11). **J**. The frequency of stimulation zone entries did not change (pre: 1.01 ± 0.04 vs stim: 1.06 ± 0.12; paired t-test, p=0.71). **K**. The latency to exit the stimulation zone significantly (pre: 0.76±0.06 vs stim: 0.26 ± 0.04; paired t-test, p< 0.0001). **L**. Sample histology of CAG-Arch injected bilaterally into the EPN. **M**. Average instantaneous speed in open field, aligned to onset of stimulation. Median (dark line) ± sem (shaded area). **N**. Inhibition of EPN had no effects on movement speed (pre: 4.87 ± 0.80 cm/s vs stim: 4.61 ± 0.47 cm/s; paired t-test, p=0.59, N=7) or **O**. real time place preference (pre: 29.56 ± 2.30% vs stim: 48.95 ± 5.51%; paired t-test, p=0.07, N=4). Mean ± sem unless otherwise noted.

The GPe is in close proximity to the entopeduncular nucleus (EPN), a basal ganglia output nucleus where inhibition could theoretically increase movement. To ensure that the GPe’s effects on speed were not caused by spillover into the EPN, we injected a separate cohort of mice with CAG-Arch specifically targeted to the EPN (Fig. 3L). Inhibiting the EPN produced no effects on locomotion (Fig. 3M-O), confirming the specificity of our previous results to inhibition of the GPe.

Unilateral inhibition of the GPe produced a turning bias in the same direction to A2A-SPN stimulation (ipsilateral), but the effect was smaller in magnitude (Fig. 3F), and full body rotations were rarely observed (Fig. 3G). In the RTPP assay, GPe inhibition was sufficient to drive robust avoidance. During the 10 minute baseline period, mice explored both sides of the arena equally (55 ± 3.2% in the inactive stimulation zone), but during the period of active stimulation, mice spent 18 ± 2.9% of their time on the stimulated side (paired t-test, p<0.0001) (Fig. 3I, *top*). This effect was similar in size to that produced by A2A-SPN stimulation (A2A-SPN Δ % time: 31.54 ± 6.99% vs CAG-Arch GPe Δ % time: 37.13 ± 3.67%; unpaired t-test, p=0.54). Once again, real-time place avoidance was driven by a shorter time spent in the stimulation zone upon entry (*latency to leave*, pre: 0.76 ± 0.06 vs stim: 0.26 ± 0.04; paired t-test, p<0.0001), rather than a decreased number of entries (*frequency of entries*, pre: 1.01 ± 0.04 vs stim: 1.06 ± 0.12; paired t-test, p=0.80) (Fig. 3J-K). Once closed loop stimulation was discontinued, mice quickly returned to exploring both sides of the arena (pre: 54.85 ± 3.22% vs post: 40.83 ± 6.87%; paired t-test, p=0.099). Mice expressing a control eYFP fluorophore, rather than Arch, showed no place preference in response to closed-loop light stimulation (Fig. 3I, *bottom*). Taken together, these results suggest that inhibition of the GPe is not sufficient to decrease movement, but is sufficient to mediate negative reinforcement.

### Behavioral effects of GPe Inhibition Are Mediated by PV-GPe Neurons

The GPe contains a heterogeneous population of neurons that can produce different effects on behavior (Cui et al., 2021a; Lilascharoen et al., 2021; Mastro et al., 2017). A recent study found that ChR2-mediated stimulation of prototypic, parvalbumin-expressing GPe neurons (PV-GPe) increases movement, whereas ChR2-mediated stimulation of a non-canonical class of GPe neuron, neuronal PAS domain protein 1-expressing GPe (Npas1-GPe), drives a decrease in movement (Cui et al., 2021a). To test whether the behavioral effects of inhibiting GPe neurons depends on which types of cell are being inhibited, we repeated our behavioral experiments, but this time, restricted Arch expression to either PV-GPe (Pvalb-2A-Cre) (Madisen et al., 2010) or Npas1-GPe (Npas1-Cre-2A-tdTomato) (Hernández et al., 2015) neuronal subpopulations.

Bilateral inhibition of PV-GPe neurons (1 mW, continuous green light, 10 trials, spaced 3 min. apart) (Fig. 4A) drove a small but significant increase in movement speed, similar to results seen with CAG-Arch (pre: 5.83 ± 0.33 cm/s vs stim: 6.90 ± 0.40 cm/s; paired t-test, p=0.002). (Fig. 4B-C). Bilateral inhibition of Npas1-GPe neurons had no consistent effect on movement speed (pre: 4.74 ± 0.43 cm/s vs stim: 5.43 ± 0.35 cm/s; paired t-test, p=0.07) (Fig. 4E-G).

**Figure 4:**
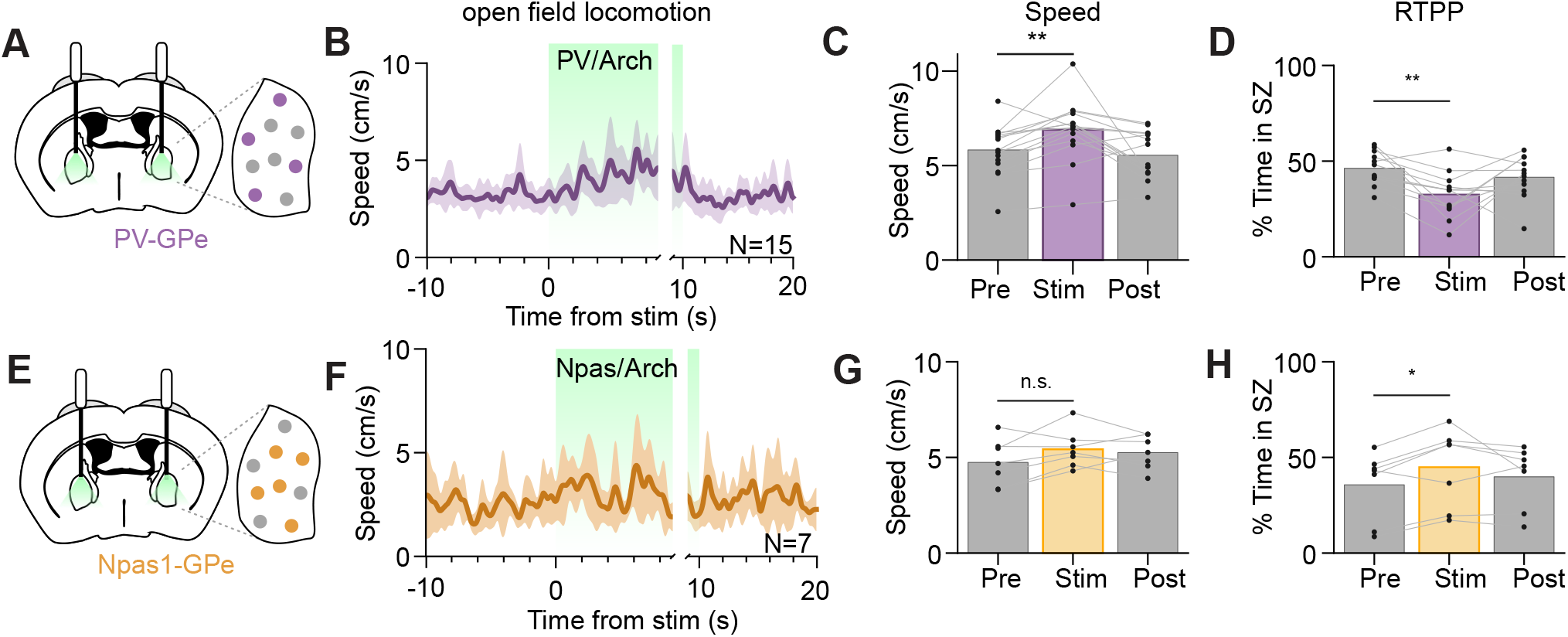
Inhibition of PV-GPe neurons is sufficient to mediate the behavioral effects of global GPe inhibition. **A**. Circuit schematic of behavioral manipulation. **B**. Instantaneous speed aligned to stimulation onset averaged across animals (N=15). Median (dark line) ± sem (shaded area). **C**. Optogenetic inhibition of PV-GPe neurons increased speed in the open field (pre: 5.83 ± 0.33 cm/s vs stim: 6.90 ± 0.40 cm/s; paired t-test, p=0.002, N=15). **D**. Bilateral optogenetic inhibition of PV-GPe neurons decreases amount of time spent in the stimulated zone (pre: 46.24 ± 2.39% vs stim: 32.55 ± 3.30%; paired t-test, p = 0.002, N=12). **E-H**. Same as top row, but for Npas1-GPe neuron inhibition. **G**. Bilateral optogenetic inhibition of Npas1-GPe neurons did not significantly increase locomotor speed (pre: 4.74±0.43 cm/s vs stim: 5.43±0.35 cm/s; paired t-test, p=0.07, N=7) **H**. Bilateral optogenetic inhibition of Npas1-GPe neurons decreased the amount of time spent in the stimulated zone (pre: 35.67 ± 6.38% vs stim: 44.89 ± 7.12%; paired t-test, p=0.02, N=7).

In the RTPP assay, inhibition of PV-GPe neurons was sufficient to drive avoidance of the stimulated side (pre: 46.24 ± 2.39% vs stim: 32.55 ± 3.30%; paired t-test, p=0.002) (Fig. 4D), which rapidly dissipated at the end of the active stimulation period (Fig. 4D). Conversely, inhibition of Npas1-GPe neurons drove a slight preference for the stimulated side (pre: 35.67 ± 6.38% vs stim: 44.89 ± 7.12%; paired t-test, p=0.02) (Fig. 4H).

These results confirm and extend our previous conclusions, that inhibition of the GPe drives negative reinforcement but not motor suppression. Inhibition of PV-GPe neurons, called “prototypic” neurons because their axons project along the main axis of the indirect pathway, drove avoidance on the RTPP assay but did not suppress movement. Inhibition of Npas1-GPe neurons, whose axons do not follow conventional indirect pathway circuitry did not affect movement, but was reinforcing in the RTPP assay.

### Optogenetic Stimulation of A2A-SPN Terminals in the GPe >1 mW Drives Antidromic Spiking and Recruits Motor Suppression

We hypothesized that if inhibition of the GPe does not dive motor suppression, perhaps this behavior is driven by activation of the inhibitory collaterals of A2A-SPNs. To test this hypothesis, we sought an approach that would preserve A2A-SPN inhibition of the GPe, but titrate the degree of collateral inhibition recruited in the striatum. To achieve this, we positioned optical fibers in the GPe to stimulate terminals of A2A-SPN neurons (A2A-Cre;Ai32) and varied the intensity of the light power to recruit different degrees of antidromic activity in the striatum (Fig. 5A).

**Figure 5:**
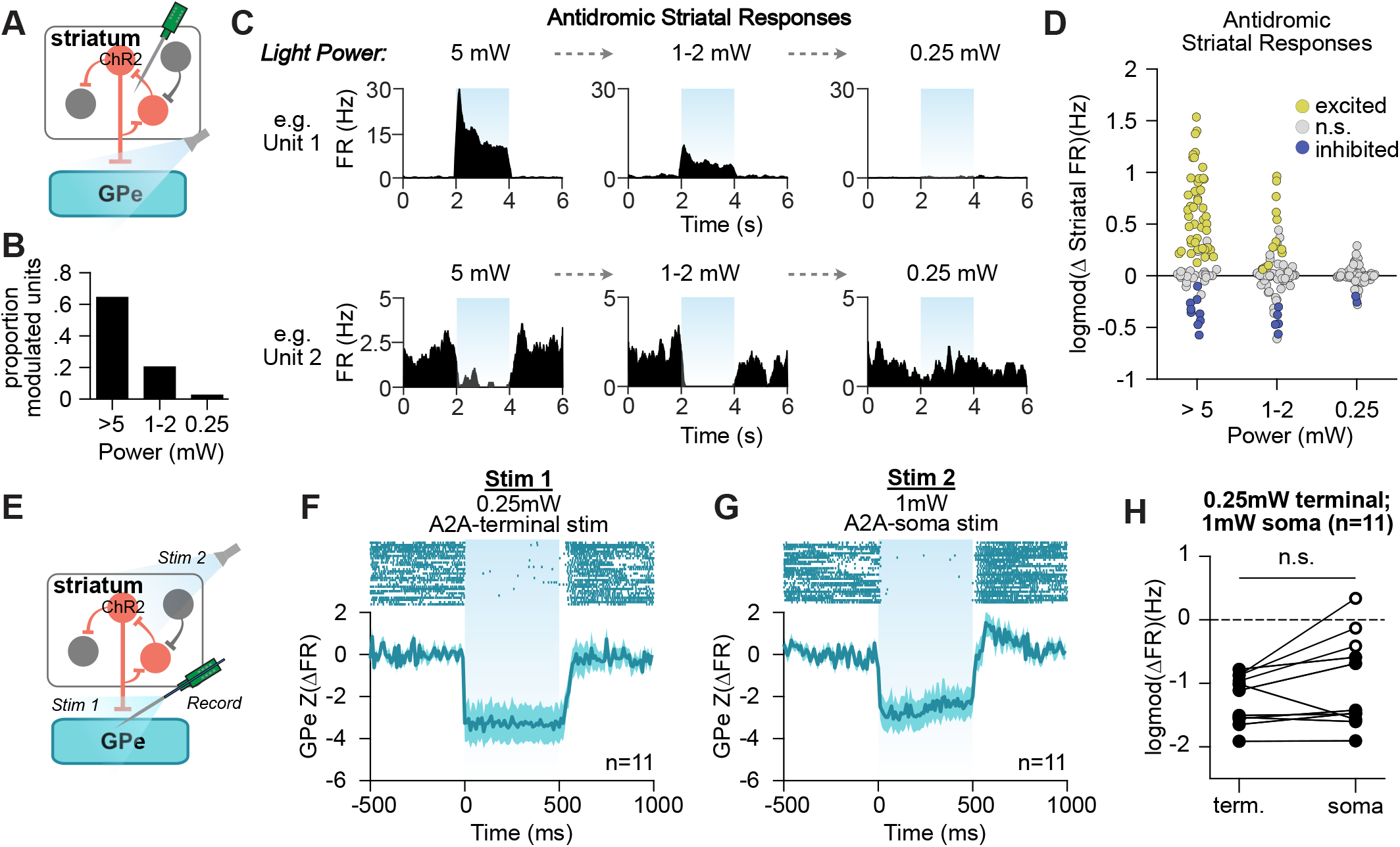
Low light A2A-terminal stimulation robustly inhibits GPe without causing antidromic activity in the striatum. **A**. Circuit schematic of *in vivo* electrophysiological recording and manipulation. **B**. Proportion of modulated units in striatum upon various stimulation light intensities of terminals in the GPe. **C**. *top:* sample excited unit recorded in the striatum across the 3 different terminal stimulation light powers (5, 1-2, and 0.25 mW). *bottom:* sample inhibited unit recorded in the striatum across the 3 different terminal stimulation light powers. **D**. Logmodulus of the change in firing rate of all striatal units (n=84). Units which were significantly excited during terminal stimulation are colored yellow. Units which were significantly inhibited during terminal stimulation are colored blue. Note that each unit recorded was held across the 3 different light stimulation powers. **E**. Circuit schematic of *in vivo* electrophysiology of GPe recordings while stimulating either A2A-terminals in GPe (0.25 mW) or A2A-somas in striatum (1 mW). **F-G**. Sample raster *(top)* and z-scored population average responses *(bottom)* of the same GPe neurons responding to 0.25 mW A2A-terminal stimulation (F) or 1 mW A2A soma stimulation (G), n = 11. **H**. Logmodulus of the change in firing rate of all GPe units for A2A-terminal (0.25mW) or A2A-soma (1mW) stimulation. GPe neurons were similarly inhibited by both manipulations (Δ-1.25 ± 0.12 Hz for A2A terminal stim and Δ-0.99 ± 0.22 Hz for A2A soma stim; Wilcoxon signed rank test; p = 0.07, n = 11). Filled circles denote significant firing rate decrease; open circles denote no significant change in firing rate.

At high light intensities (5 – 10 mW), stimulation of A2A-SPN terminals in the GPe drove firing rate changes in 54/84 of striatal units (64.2%, across 4 mice) (Fig. 5B), of which 45 were excited (Δ0.65 ± 0.06 Hz) and 9 were inhibited (Δ-0.35 ± 0.05 Hz) (Fig. 5C-D). Lowering stimulus intensity to 2 mW reduced but did not eliminate antidromic activity within the striatum; 19/84 units (22.6%) were significantly modulated (Fig. 5B), with 14 showing excitation (Δ0.42 ± 0.08 Hz) and 9 showing inhibition (Δ-0.44 ± 0.04 Hz) (Fig. 5C-D). Further reducing light intensity to 0.25 mW eliminated nearly all antidromic activity within the striatum (Fig. 5B). Only 2/84 striatal units were significantly inhibited during stimulation (Δ-0.42 ± 0.05 Hz on average) (Fig. 5C-D).

To ensure that 0.25 mW stimulation of A2A-SPN terminals was still sufficient to inhibit the GPe, we performed recordings from mice in which optrodes were simultaneously positioned in both the striatum and GPe (Fig. 5E). As shown in Fig. 5F-H, individual GPe neurons were similarly inhibited by 0.25 mW stimulation of A2A-SPN terminals as they were by 1 mW stimulation of A2A-SPN somas (as used in previous behavioral experiments, Fig. 1) (Δ-1.25 ± 0.12 Hz for A2A-terminal stim and Δ-0.99 ± 0.22 Hz for A2A-soma stim; Wilcoxon signed rank test; p= 0.07, n=11).

The effects of increasing antidromic recruitment of the striatal network on movement was assessed in the open field by stimulating A2A-SPN terminals in the GPe of A2A-Cre; Ai32 mice (Fig. 6A). Bilateral optical fibers were implanted into the GPe and 2 s light pulses of varying intensity (0.1 – 3 mW) were delivered at random inter-stimulus intervals of 7-14 s for a total of 50 trials per mouse. Effects on locomotor speed were calculated by comparing the average speed during 2 s of stimulation to the average speed 2 s before stimulation (speed_during_ – speed_before_). Fig. 6B shows the effects on movement speed across the various light intensities tested (N = 5 mice). At low light levels, stimulation of A2A-SPN terminals in the GPe had no consistent effect on movement speed (0-1 mW, Fig. 6B-C). But as the power of optical stimulation was increased, motor effects began to emerge (Fig. 6B,C), becoming significant at light powers > 1.61 ± 0.2 mW. As light power was further increased, motor suppression became even more pronounced, plateauing between 2-3 mW (Fig. 6B,C) (Δspeed_baseline_: 0.89 ± 0.41 cm/s vs Δspeed_stim_: -2.22 ± 0.38 cm/s at 1.61 mW; paired t-test, p=0.045). Together with the results from Fig. 5, these data suggest that the motor effects of stimulating A2A-SPN terminals in the GPe begin to appear at the same stimulus intensities that recruit antidromic activity and collateral inhibition within the striatum.

**Figure 6:**
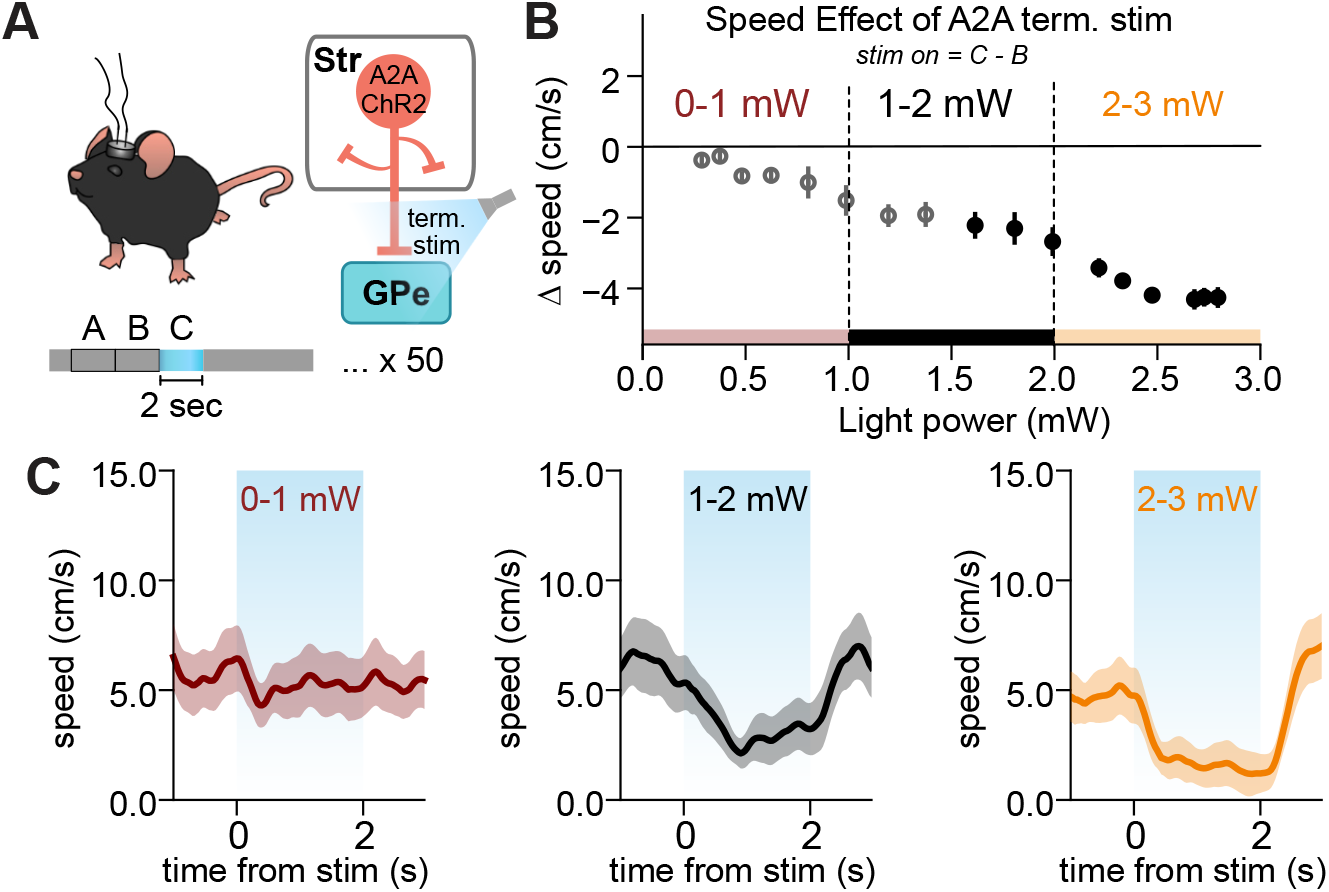
Movement suppression is correlated with the light power of A2A-terminal stimulation. **A**. Circuit schematic of behavioral manipulation. Mice received fifty, 2 s bilateral stimulations of A2A-terminals in the GPe. Each stimulation was a random light power ranging from 0-3 mW achieved by pulse-width modulation. **B**. Change in movement speed (average speed 2 seconds prior to stimulation subtracted from average speed during stimulation period) as a function of average light power. Closed black circles denote significant changes in speed compared to pre-stim movement periods (at 1.61mW: Δspeed_baseline(B-A)_=0.89 ± 0.41 cm/s vs Δspeed_stim(C-B)_=-2.22 ± 0.38 cm/s at 1.61 mW; paired t-test, p=0.045, N = 5 mice). **C**. Average movement speed aligned to the onset of stimulation, grouped into trials where light power was 0-1 mW, 2-3 mW, or 2-3 mW.

To test the hypothesis that A2A-SPN projections to the GPe mediate RTPP avoidance, whereas their collaterals in the striatum mediate motor suppression, we repeated open field behavioral experiments but this time stimulated terminals of A2A-SPNs in the GPe at 0.25 mW or 2 mW (Fig. 7A,C). Stimulation at 0.25 mW is expected to inhibit the GPe, but not drive antidromic activity in the striatum; stimulation at 2 mW is expected to inhibit the GPe *and* recruit antidromic activity in the striatum.

**Figure 7:**
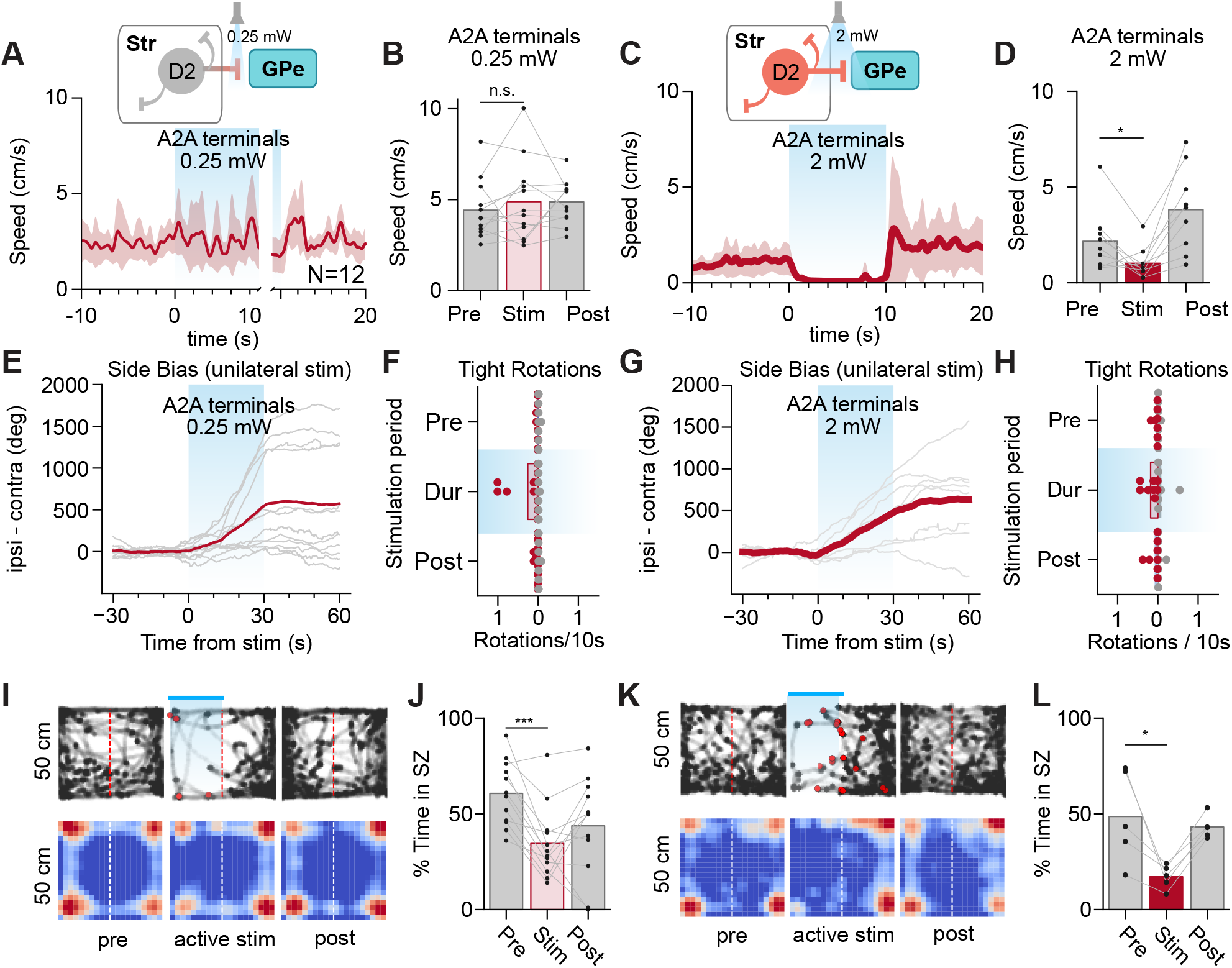
A2A-SPN projections to the GPe mediate RTPP avoidance, but not motor suppression. **A**. Low light power (0.25mW) A2A-terminal stimulation in the GPe does not affect movement speed in the open field. **B**. Average speed during stimulation epoch is not different from pre-stimulation speed (pre: 4.42 ± 0.44 cm/s vs stim: 4.89 ± 0.62; paired t-test, p=0.37, N=12). **C**. High light power (2mW) A2A-terminal stimulation in the GPe causes a speed slow-down in the same mice. **D**. Average speed during 2mW stimulation of A2A-terminals in the GPe is significantly slower compared to pre-stimulation periods (pre: 2.17 ± 0.52 cm/s vs stim: 1.02 ± 0.26 cm/s; paired t-test, p=0.027). **E**,**G**. Unilateral stimulation of A2A-terminals at 0.25mW and 2 mW elicits an ipsilateral turn bias. **F**,**H**. Unilateral stimulation of A2A-terminals at 0.25 mW and 2mW does not drive tight ipsilateral rotations. **I**. 0.25mW A2A-terminals stimulation drives robust avoidance in a RTPP task. **J**. Percent time spent in stimulated zone significantly decreases during active stimulation period compared to baseline (pre: 60.68 ± 4.68% vs stim: 34.54± 5.32%; paired t-test; p=0.0002). **K**. 2mW A2A-terminal stimulation drives robust avoidance and more freezing/slow-down bouts in a RTPP task. **L**. Percent time spent in stimulated zone significantly decreases during the active stimulation period compared to baseline (pre: 48.62 ± 9.65% vs stim: 17.15 ± 2.50%; paired t-test, p=0.041).

When A2A-SPN terminals were stimulated at 0.25 mW, mice showed no consistent changes in locomotor speed (Fig. 7A-B), but did show significant slowing of movement speed when terminals were stimulated at 2 mW (speed change, 0.25mW vs 2mW: Δ +0.46 ± 0.48 cm/s vs. Δ -1.15 ± 0.40 cm/s; paired t-test, p=0.03) (Fig. 7C-D). Conversely, both stimulus intensities were equally effective at driving avoidance on the RTPP task (% change in stim zone time, 0.25 mW vs. 2 mW: Δ -26.14 ± 4.61% vs Δ -31.47 ± 9.47%; paired t-test, p=0.60) (Fig. 7I-L). Unilateral stimulation at both powers produced similar effects on side bias, measured as accumulated angular displacement (Fig. 7E,G), but few full body rotations (Fig. 7F,H).

Taken together, these results demonstrate that A2A-SPN-mediated inhibition of the GPe is sufficient to drive avoidance on the RTPP assay, but does not decrease movement speed. Movement speed is only decreased when stimulation of A2A-SPN terminals is sufficiently strong that it recruits antidromic activity within the striatum.

## Discussion

Optogenetic stimulation of A2A-SPNs is a common experimental approach used to study the behavioral correlates of the indirect pathway. A primary function of the indirect pathway is thought to be motor suppression, reinforced by the fact that optogenetic stimulation of A2A-SPNs inhibits movement (Bakhurin et al., 2020; Kravitz et al., 2010; Roseberry et al., 2016). Our results challenge this assumption by demonstrating that the motor suppressing effects of A2A-SPN stimulation are not driven by inhibition of the GPe and the indirect pathway, but rather through collateral inhibition in the striatum.

### Behavioral Correlates of the Indirect Pathway

The hypothesis that the indirect pathway is a motor suppression pathway was formulated based on anatomical and clinical studies (Albin et al., 1989; DeLong, 1990), and has been highly influential in basal ganglia research. The indirect pathway is complex, spanning connections across multiple basal ganglia nuclei (Grillner and Robertson, 2016; Kita, 2007; Smith et al., 1998). Within the striatum, neurons projecting to the indirect pathway are molecularly distinct from those projecting to the direct pathway (Gerfen et al., 1990), leading to current conventions where the activity of Drd1-SPNs is taken as synonymous with direct pathway activity, and activity of A2A-SPNs is taken as synonymous with indirect pathway activity. Kravitz et al. demonstrated that driving Drd1-SPNs facilitated movement, while driving Drd2-SPNs (homologous to A2A-SPNs) inhibited movement, a result widely accepted as evidence of the opposing effect of the direct and indirect pathways on locomotion (Kravitz et al., 2010). However, subsequent studies of SPN activity during spontaneous behaviors found that Drd1 and A2A SPNs were often co-activated during movement (Cui et al., 2013; Markowitz et al., 2018), consistent with an action selection model, rather than a go/stop model (Jin et al., 2014; Mink, 1996; Sales-Carbonell et al., 2018).

In the action selection framework, the direct pathway carries information about actions to be executed, while the indirect pathway filters out competing actions to be suppressed (DeLong and Wichmann, 2009; Gerfen and Surmeier, 2011; Mink, 1996). As animals learn to predict which actions in a given environment are likely to lead to rewards or punishments, they update their actions accordingly, a process called reinforcement learning (Dayan and Niv, 2008). Optogenetic stimulation of Drd1-SPNs is reinforcing, meaning it maintains or increases execution of a given behavior, while optogenetic stimulation of A2A-SPNs is punishing, meaning it decreases execution of a given behavior (Kravitz et al., 2012; Lalive et al., 2018; Matamales et al., 2020; Yttri and Dudman, 2016). Although the motor and reinforcing effects of the direct and indirect pathways are often studied independently, attributed to parallel but segregated motor/limbic loops within the basal ganglia (Alexander et al., 1986), these behavioral features are more integrated than typically assumed. The indirect pathway has little impact on movement in the absence of decision making (Bolkan et al., 2022), and animals can be trained such that stimulation of A2A-SPNs drives increases in movement velocity, rather that decreases (Yttri and Dudman, 2016).

### Motor Functions of the GPe

In the context of ongoing debates about the role of the direct and indirect pathways on behavior, comparatively few studies have looked outside the striatum at neural activity in downstream basal ganglia nuclei. The GPe is the sole output target of A2A-SPNs, but firing rates of GPe neurons, in aggregate, are poor predictors of movement velocity (Dodson et al., 2015; Gu et al., 2020; Kimura et al., 1996; Mink and Thach, 1991; Turner and Anderson, 1997). Most GPe neurons exhibit motor-related activity, but some neurons increase firing rates during movement while others decrease firing rates, and there is little evidence to suggest that the GPe is broadly inhibited during stopping, as predicted by the classic rate model of the indirect pathway. A recent study suggested that GPe neurons are involved in proactive inhibition, a predictive form of behavioral control in which GPe neurons help to slow reaction times under conditions where stop cues might appear (Gu et al., 2020). This would suggest that GPe neurons carry information about environmental or behavioral contexts. Indeed, analyses of GPe firing rates during complex tasks have revealed that individual GPe neurons independently encode information about both movement and value (Arkadir et al., 2004; Kim et al., 2017) and on a reversal learning task GPe neurons provide a reinforcement signal that predicts task performance (Schechtman et al., 2016). Our results support a prominent role for the GPe in behavioral punishment, but not the slowing of movement velocity. Inhibition of the GPe, either directly with Arch, or synaptically by A2A-SPNs, did not reduce movement speed, but did impact where mice directed their actions, causing them to spend less time in areas of the arena that drove a decrease in GPe activity.

A previous study found that stimulation of A2A-SPN terminals in the GPe resulted in motor suppression (Cazorla et al., 2014), but results from our study suggest that this effect might have been driven by off-target, antidromic activation of the striatum. Other studies have reported motor effects when GPe neurons are optogenetically excited (Aristieta et al., 2021; Cui et al., 2021a; Lilascharoen et al., 2021). The major excitatory input to the GPe comes from the subthalamic nucleus (STN), which is part of the hyperdirect pathway of the basal ganglia that drives rapid behavioral stopping (Dunovan et al., 2015; Guillaumin et al., 2021; Nambu et al., 2002; Polyakova et al., 2020). It is not known whether the GPe is part of the neural circuitry required for the stopping effects of the hyperdirect pathway.

Optogenetic stimulation of an atypical class of neurons in the GPe, arkypallidal neurons, drives robust suppression of movement (Aristieta et al., 2021; Cui et al., 2021a). Arkpallidal neurons make up ∼25% of GPe neurons and project exclusively back to the striatum in a broad, “net-like” pattern, distinguishing them from prototypical GPe neurons whose downstream projections contribute to the indirect pathway (Mallet et al., 2012). Widespread inhibition of the striatum by arkypallidal neurons is thought to provide a cancelling signal that aborts upcoming or ongoing actions (Mallet et al., 2016). A recent study suggested that during indirect pathway signaling, arkypallidal GPe neurons are disinhibited, providing a potential mechanism by which indirect pathway signaling could drive motor suppression (Aristieta et al., 2021). But our results found that under conditions in which inhibition from A2A-SPNs was restricted to the GPe (terminal stimulation experiments), motor suppression was not observed. Taken together, these results suggest the presence of a neural circuit running through the GPe that can suppress movement, but this circuit is distinct from the canonical indirect pathway.

Both somatic stimulation of A2A-SPNs and stimulation of arkypallidal GPe neurons drive motor suppression, and share a common mechanism of driving widespread inhibition in the striatum (in the case of A2A-SPNs, through inhibitory collaterals) (Burke et al., 2017; Dobbs et al., 2016; Taverna et al., 2008). In the ventral striatum, collaterals of A2A-SPNs regulate the degree of locomotor sensitization observed following cocaine administration (Dobbs et al., 2016). Collateral inhibition of Drd1-SPNs in the direct pathway would be predicted to reduce movement, and indeed, this was the result when Drd1-SPNs were optogenetically inhibited (Tecuapetla et al., 2016). However, the degree of motor suppression was weaker than what has been observed during A2A-SPN stimulation, which could reflect technical limitations of inhibiting an already sparsely firing population, or could suggest the presence of motor suppressing ensembles in the striatum that have not yet been fully characterized (Cui et al., 2021b; Parker et al., 2018).

### Implications for Disease

Motor deficits of Parkinson’s disease (PD) have been hypothesized to reflect overactivity of the indirect pathway: increased firing of A2A-SPNs and excessive inhibition of the GPe (Albin et al., 1989; DeLong, 1990). It is possible that the indirect pathway takes on a greater role in motor control in disease, when changes in indirect pathway signaling are more chronic. Sustained inhibition of the GPe using DREADDs reduced movement by ∼30% (Lilascharoen et al., 2021). But data collected from the striatum of human PD patients found no evidence of increased firing rates in A2A-SPNs (Valsky et al., 2020), and the loss of striatal dopamine was recently shown to have a bigger impact on learning, rather than motor deficits (González-Rodríguez et al., 2021). In dopamine depleted mice, input from A2A-SPNs contributes to the excessive patterning of GPe neurons that is a physiological hallmark of the parkinsonian state (Kovaleski et al., 2020). Both the GPe and its reciprocally connected partner, the STN, are effective targets for alleviating motor symptoms in PD (Mastro et al., 2017; Schor and Nelson, 2019; Spix et al., 2021; Wichmann and DeLong, 2011; Yu et al., 2020), but the therapeutic efficacy might have more to do with disrupting aberrant firing patterns, rather than changing the level of indirect pathway signaling (Bevan et al., 2002; Eusebio et al., 2012; Nelson and Kreitzer, 2014).

### Conclusions

In conclusion, our results suggest that under normal conditions, the indirect pathway plays a greater role in shaping behavior through punishment, rather than regulating gross locomotor function. Our results emphasize the importance of understanding the effects of optogenetic manipulations on broader network activity, not just the cell populations being stimulated, and challenge assumptions that firing rates of A2A-SPNs are synonymous with indirect pathway activity.

## Methods

### RESOURCE AVAILABILITY

#### Lead contact

Further information and requests for resources and reagents should be directed to and will be fulfilled by the Lead Contact, Aryn Gittis.

#### Materials availability

This study did not generate new unique reagents.

#### Data and code availability

- All data reported in this paper will be shared by the lead contact upon request.
- Custom MATLAB and Python scripts developed by our laboratory to analyze mouse behavior and electrophysiology data are available upon request.
- Any additional information required to reanalyze the data reported in this paper is available from the lead contact upon request.

### EXPERIMENTAL MODEL AND SUBJECT DETAILS

#### Animals and surgery

Experiments were conducted in accordance with the guidelines from the National Institutes of Health and with approval from Carnegie Mellon University Institutional Animal Care and Use Committee. Adult male and female mice (>8 weeks old) on a C57BL/6J background were used for experiments. All mice were maintained on a reverse circadian 12-h light/12-h dark cycle with food and water provided *ad libitum*. All surgical procedures were performed using aseptic techniques and were performed using general anesthesia. Post-surgery, animals were injected with 3mg/kg Ketofen s.c. and allowed to recover on a heating pad before being returned to their standard home cage.

#### AAV vectors

Injections of purified double-floxed AAV2-DIO-eYFP (controls), AAV2-DIO-ChR2-eYFP (cell-specific activation), AAV2-DIO-Arch-eYFP (cell-specific inhibition), or AAV2-CAG-Arch-GFP (nonspecific inhibition) (Addgene) were made in A2A-Cre, PV-Cre, or Npas-Cre transgenic mice 8–12 weeks old.

#### Stereotaxic injections

AAV vectors were injected into the dorsomedial striatum (DMS) or globus pallidus externus (GPe) and completed in accordance with methods previously described (Mastro et al., 2014). Briefly, anesthesia was induced using ketamine (100 mg/kg) and xylazine (30 mg/kg) and maintained throughout surgery using 1.5% isoflurane. Mice were placed in a stereotaxic frame (Kopf Instruments), where the scalp was opened and bilateral holes were drilled in the skull (striatum: 1-2 mm AP, +/-1.5 mm ML, GPe: 0.1 mm AP, +/-2.2 mm ML from Bregma). Virus (Striatum: 0.5-1 μl, GPe: 150 nL) was injected with a Nanoject (Drummond Scientific) through a pulled glass pipet (tip diameter ∼30 μm) whose tip was positioned below the top of the skull (striatum: -2.5 mm DV, GPe: -3.68 mm DV). To prevent backflow of virus, the pipet was left in the brain for 5 min after completion of the injection. During the same surgery, optic fiber cannulae were implanted using dental cement (see: optogenetic simulation). All experiments were performed at least 2 weeks after injection to allow time for full viral expression. At that point, some mice underwent a second surgery to make craniotomies for electrophysiology.

#### Optogenetic stimulation

For all implants, optic fibers (Thorlabs), cut to length and glued inside 1.25mm ceramic ferrules, were matched for light transmission and calibrated to achieve the desired output measured at the tip of optic fiber cannula (PM100D, Thorlabs). Inhibition of GPe was performed with constant 1mW green light from high powered LEDs (545nm, UHP-T-545-SR, Prizmatix). Excitation of Str was performed with constant 1mW blue light from high powered LEDs (450 nm, UHP-T-450-SR, Prizmatix). During optogenetic light tritration curves and *in vivo* iSPN terminal stimulation, a blue laser was used with pulse-width-modulation to deliver intensities from 0-10mW (473nm, LRS-0473-GFM-00100-05, Laserglow Technologies).

#### *In vivo* recordings

Head-bar implants to secure mice for *in vivo* recordings were performed under anesthesia asdescribed above. Bilateral craniotomies (for probe insertion) were created over striatum or GPe using coordinates described above and a stainless steel headbar was implanted using Metabond dental cement. Dental cement was extended from the head-bar to surround the extent of both craniotomies to form a well. This well was then filled with silicone elastomer (Kwik-sil, WPI) that prevented infection and damage to the exposed brain tissue and the mouse was allowed to recover for 24 hours. During the recording, this well was filled with 0.9% NaCl and used as a ground reference. On the day of recording, animals were fixed to the top of the wheel and allowed 15 min to acclimate to the head-fixed position. The silicone elastomer was removed and a linear 16-channel silicon probe was attached to the micromanipulator and centered on Bregma. The probe was slowly advanced (5–7um/s) until the top of the recording target was reached (Str: -2-3mm DV, GPe: -3 DV from the top of the brain). Post-mortem tissue analysis of electrode tracks and Iba-1 (Wako) immunoreactivity induced by probe penetrations were further evidence of proper targeting. Once a population of responsive units was identified, optogenetic stimulation was performed (see: optogenetic stimulation).

#### Histological procedures

Mice were anesthetized with a lethal dose of ketamine (100 mg/kg) and xylazine (30 mg/kg) and perfused transcardially with phosphate buffered saline (PBS), followed by 4% paraformaldehyde (PFA) in PBS. Brains were retrieved and fixed in 4% PFA for 24-hours before being placed in a 30% sucrose solution until sinking. Whole brains were sectioned in 30 μm slices using a freezing microtome (Microm HM 430; Thermo Scientific). Sections were scanned with a fluorescence microscope (BZ-X series, Keyence).

#### Iba-1 immunoreactivity

Recording probe location was visualized with immunofluorescence against the microglial marker, Iba-1 (rabbit anti-Iba-1, 1:1000, Wako). Primary antibodies were detected with Alexa Fluor 647-conjugated donkey anti-rabbit (1:500, Thermo Fisher Life Technologies), incubated for 90 min at room temperature, or Alexa Fluor 488 donkey anti-rabbit (1:500, Thermo Fisher Life Technologies), incubated for 3 hours at room temperature.

### QUANTIFICATION AND STATISTICAL ANALYSIS

Statistical details for each experiment can be found in the corresponding figure legends. All data are expressed as means and standard errors of the mean (SEM) unless otherwise noted. We assessed the statistical significance using parametric (t-tests and ANOVA) and nonparametric (Wilcoxon) tests; significance was considered for test statistics with a (*) p-value of < 0.05.

#### Behavioral optogenetic assessment

Animals received optogenetic stimulation in the following contexts: bilaterally and unilaterally in open field and in a real-time place preference task. The minimum interval between two consecutive behavioral assays was typically 12 hours. Mice were habituated to the testing room for 20 min before testing and the arena was cleaned with 50% ethanol in between animals. Mouse behavior was captured by top-down and side cameras (29.97 FPS). Positions of nose, tail, and center of mass of each mouse were tracked using EthoVision XT software (Noldus) and refined using DeepLabCut for assessing rotations. Distance traveled and average speeds for the data acquisition period were calculated using EthoVision and custom software. Ambulation bouts were scored post hoc as periods of >2cm/s movement lasting for > 0.5s and >0.5s apart) (Kravitz et al., 2010). Immobility bouts were scored by EthoVision using a criterion of <2% pixel change in 0.5s, lasting for >0.5s and >0.5s apart. Fine movement bouts were scored as periods where the mouse was not ambulating and not immobile lasting for >0.5s and >0.5s apart.

#### Open field optogenetics

Mice were placed in a ∼50 cm^2^ clear square open field and experienced a 10 m habituation period. After this, mice received 10 stimulations of 10-30 s in duration (see: behavioral optogenetic stimulation) spaced 2-3 minutes apart. Behavioral effects of stimulation were assessed in the first 10s of stimulation and compared to baseline period (10 s prior to stimulation onset).

#### Optogenetic real-time place preference

The same open field arena was divided into two zones. After a 10 m baseline period, a 10-30 m active stimulation period began with the tracked mouse body center triggering optogenetic stimulation in a closed-loop in one of the two zones. If the mouse remained in the active zone for longer than 30 s, the stimulation reverted to a 2 s on 8 s off pattern until the mouse exited the zone. This allowed mice an opportunity to leave the zone if stimulation caused complete motor arrest. The active period was followed by at 10-30 m post period with no stimulation. Only the first 10 m were assessed of active and post periods. Place preference was assessed by examining the percent time spent in the stimulated zone during the active portion of the task relative to baseline period (no stimulation). The overall density of open field occupancy was assessed by generating a 2D histogram of time spent in 2.5cm^2^ bins of the arena and expressed as a probability density function in Python (number of samples in bin / number samples / bin area; numpy.histogram2d(density=True)). These plots were then smoothed with a 2D gaussian kernel and averaged across mice for visualization purposes. The number and duration of stimulation zone entrances were assessed throughout the experiment in 1 minute bins and normalized to the average baseline period value for each mouse.

#### Rotations

Mice were placed in the same open field and the light source was delivered to either the left or right optic fiber cannula only. Five 30 s stimulations were applied, 180 s apart and rotations were assessed relative to the 30 s baseline period prior to stimulation. DeepLabCut was then used to track mouse nose and tail tip from a top-down camera. Mouse body angle was calculated from these coordinates. A rotation was scored when a mouse completed a 360º turn within 10 s. A minimum criterion of 0.3 s was used to exclude artifactual turning. Cumulative degree change was also assessed (ipsilateral – contralateral to stimulation location) to assess more subtle bias in mouse movement direction.

#### Optogenetic titration curve

To assess the effect of various light intensities on locomotion when stimulating iSPN terminals in GPe, mice experienced 50, 2 s bilateral stimulations with randomized light intensity from 0 to ∼3 mW achieved by pulse-width modulation (PWM) of a blue laser (473nm) using custom hardware (Arduino Uno). Stimulations were delivered at random inter-stimulus intervals of 7-14 s using Ethovision as described above to control trial structure and video monitoring. Fiber cannulas were calibrated at the tip for 0-10 mW output using values 0-255 of Arduino PWM output to reconstruct stimulation intensity.

#### Spike sorting

Data was sampled at 32 kHz and filtered at 150–8000 Hz for spiking activity using Neuronexus 16 channel linear probes with site spacing of 25-50um (A1×16-5mm-50-177-A16, A1×16-10mm-25-177-A16, and A1×16-5mm-50-177-OA16LP 0.66 numerical aperture, Neuronexus). All spike detection, clustering and sorting was performed using UltraMegaSort 2000. Linear probe electrode sites were broken into groups of 4-adjacent sites (tetrodes) and spike events were detected 4 SD below the mean with a 0.75 ms shadow period. Events were then clustered using k-means clustering (k-means cluster size = 50), aggregated based on interface energy (agg_cutoff = 0.2; Fee et al., 1996) and manually inspected on the basis of spike waveform, stability across time, and inter-spike interval refractory period violations (< 0.5% of intervals < 1.5ms).

#### Neural data analysis

Neural responses to optogenetic stimulation were evaluated by comparing the firing rate preceding and following stimulus onset using a paired t-test with alpha = 0.05. PSTHs were calculated from 10-100 stimulus trials and smoothed with a gaussian convolution. Z-scored PSTHs used the mean and standard deviation of baseline firing to normalize stimulus response. The log-modulus transformation (L(x)=sign(x)*log(|x|+1)) was performed on recorded striatal units to appropriately assess inhibitory and excitatory responses upon stimulation. Striatal optotagging was performed by identifying neurons with an excitatory response that occurred within 15ms of stimulus onset (0.25-1 mW, paired t-test, alpha = 0.01, 200-300 trials). Tagged units were required to exhibit tagged response at the beginning and end of recordings to be considered tagged. Striatal inhibited units were required to show a significant inhibitory response to direct striatal stimulation within 200ms (paired t-test, alpha = 0.01). Antidromic modulation of striatal units by stimulation of terminals in GPe was evaluated using 2s long stimulations at low (0.25 mW), high (∼1-2 mW), or very high (5-10 mW) constant light (30-50 trials). GPe inhibition by A2A terminals was evaluated using an optrode inserted into GPe and 1mW constant light at 0.5 to 10 s. GPe inhibition by CAG Arch was evaluated using green light through optrode in GPe (1mW).

## Notes

### Competing Interest Statement

The authors have declared no competing interest.

## References

Albin, R.L., Young, A.B., and Penney, J.B. (1989). The functional anatomy of basal ganglia disorders. Trends Neurosci 12, 366–375.

Alexander, G.E., DeLong, M.R., and Strick, P.L. (1986). Parallel organization of functionally segregated circuits linking basal ganglia and cortex. Annu Rev Neurosci 9, 357–381.

Aristieta, A., Barresi, M., Lindi, S.A., Barriere, G., Courtand, G., de la Crompe, B., Guilhemsang, L., Gauthier, S., Fioramonti, S., and Baufreton, J. (2021). A disynaptic circuit in the globus pallidus controls locomotion inhibition. Curr. Biol. 31, 707–721.

Arkadir, D., Morris, G., Vaadia, E., and Bergman, H. (2004). Independent coding of movement direction and reward prediction by single pallidal neurons. J. Neurosci. 24, 10047–10056.

Bakhurin, K.I., Li, X., Friedman, A.D., Lusk, N.A., Watson, G.D., Kim, N., and Yin, H.H. (2020). Opponent regulation of action performance and timing by striatonigral and striatopallidal pathways. Elife 9.

Bevan, M.D., Magill, P.J., Terman, D., Bolam, J.P., and Wilson, C.J. (2002). Move to the rhythm: oscillations in the subthalamic nucleus-external globus pallidus network. Trends Neurosci 25, 525–531.

Bolkan, S.S., Stone, I.R., Pinto, L., Ashwood, Z.C., Iravedra Garcia, J.M., Herman, A.L., Singh, P., Bandi, A., Cox, J., and Zimmerman, C.A. (2022). Opponent control of behavior by dorsomedial striatal pathways depends on task demands and internal state. Nat. Neurosci. 25, 345–357.

Burke, D.A., Rotstein, H.G., and Alvarez, V.A. (2017). Striatal Local Circuitry: A New Framework for Lateral Inhibition. Neuron 96, 267–284.

Cazorla, M., Delmondes De Carvalho, F., Chohan, M.O., Shegda, M., Chuhma, N., Rayport, S., Ahmari, S.E., Moore, H., and Kellendonk, C. (2014). Article Dopamine D2 Receptors Regulate the Anatomical and Functional Balance of Basal Ganglia Circuitry. Neuron 81, 153–164.

Cox, J., and Witten, I.B. (2019). Striatal circuits for reward learning and decision-making. Nat. Rev. Neurosci. 20, 482–494.

Cui, G., Jun, S.B., Jin, X., Pham, M.D., Vogel, S.S., Lovinger, D.M., and Costa, R.M. (2013). Concurrent activation of striatal direct and indirect pathways during action initiation. Nature 494, 238–242.

Cui, Q., Pamukcu, A., Cherian, S., Chang, I.Y.M., Berceau, B.L., Xenias, H.S., Higgs, M.H., Rajamanickam, S., Chen, Y., Du, X., et al. (2021a). Dissociable Roles of Pallidal Neuron Subtypes in Regulating Motor Patterns. J. Neurosci. 41, 4036–4059.

Cui, Q., Du, X., Chang, I.Y.M., Pamukcu, A., Lilascharoen, V., Berceau, B.L., García, D., Hong, D., Chon, U., Narayanan, A., et al. (2021b). Striatal Direct Pathway Targets Npas1 + Pallidal Neurons. J. Neurosci. JN-RM-2306-20.

Dayan, P., and Niv, Y. (2008). Reinforcement learning: the good, the bad and the ugly. Curr. Opin. Neurobiol. 18, 185–196.

DeLong, M.R. (1990). Primate models of movement disorders of basal ganglia origin. Trends Neurosci. 13, 281–285.

DeLong, M., and Wichmann, T. (2009). Update on models of basal ganglia function and dysfunction. Parkinsonism Relat. Disord. 15, S237–S240.

Dobbs, L.K., Kaplan, A.R., Lemos, J.C., Matsui, A., Rubinstein, M., and Alvarez, V.A. (2016). Dopamine Regulation of Lateral Inhibition between Striatal Neurons Gates the Stimulant Actions of Cocaine. Neuron 90, 1100–1113.

Dodson, P.D., Larvin, J.T., Duffell, J.M., Garas, F.N., Doig, N.M., Kessaris, N., Duguid, I.C., Bogacz, R., Butt, S.J.B., and Magill, P.J. (2015). Distinct developmental origins manifest in the specialized encoding of movement by adult neurons of the external globus pallidus. Neuron 86, 501–513.

Dunovan, K., Lynch, B., Molesworth, T., and Verstynen, T. (2015). Competing basal ganglia pathways determine the difference between stopping and deciding not to go. Elife 4, e08723.

Eusebio, A., Cagnan, H., and Brown, P. (2012). Does suppression of oscillatory synchronisation mediate some of the therapeutic effects of DBS in patients with Parkinson’s disease? Front. Integr. Neurosci. 6, 47.

Freeze, B.S., Kravitz, A. V, Hammack, N., Berke, J.D., and Kreitzer, A.C. (2013). Control of Basal Ganglia Output by Direct and Indirect Pathway Projection Neurons. J. Neurosci. 33, 18531–18539.

Gerfen, C.R., and Surmeier, D.J. (2011). Modulation of striatal projection systems by dopamine. Annu. Rev. Neurosci. 34, 441–466.

Gerfen, C.R., Engber, T.M., Mahan, L.C., Susel, Z., Chase, T.N., Monsma Jr., F.J., and Sibley, D.R. (1990). D1 and D2 dopamine receptor-regulated gene expression of striatonigral and striatopallidal neurons. Science (80-.). 250, 1429–1432.

Gong, S., Doughty, M., Harbaugh, C.R., Cummins, A., Hatten, M.E., Heintz, N., and Gerfen, C.R. (2007). Targeting Cre recombinase to specific neuron populations with bacterial artificial chromosome constructs. J. Neurosci. 27, 9817–9823.

González-Rodríguez, P., Zampese, E., Stout, K.A., Guzman, J.N., Ilijic, E., Yang, B., Tkatch, T., Stavarache, M.A., Wokosin, D.L., Gao, L., et al. (2021). Disruption of mitochondrial complex I induces progressive parkinsonism. Nature 599, 650–656.

Grillner, S., and Robertson, B. (2016). The Basal Ganglia Over 500 Million Years. Curr. Biol. 26, R1088–R1100.

Gu, B.-M., Schmidt, R., and Berke, J.D. (2020). Globus pallidus dynamics reveal covert strategies for behavioral inhibition. Elife 9.

Guillaumin, A., Serra G. Pietro, Georges, F., and Wallén-Mackenzie, Å. (2021). Experimental investigation into the role of the subthalamic nucleus (STN) in motor control using optogenetics in mice. Brain Res. 1755, 147226.

Hernández, V.M., Hegeman, D.J., Cui, Q., Kelver, D.A., Fiske, M.P., Glajch, K.E., Pitt, J.E., Huang, T.Y., Justice, N.J., and Chan, C.S. (2015). Parvalbumin<sup>+</sup> Neurons and Npas1<sup>+</sup> Neurons Are Distinct Neuron Classes in the Mouse External Globus Pallidus. J. Neurosci. 35, 11830 LP – 11847.

Horak, F.B., and Anderson, M.E. (1984). Influence of globus pallidus on arm movements in monkeys. II. Effects of stimulation. J Neurophysiol 52, 305–322.

Jin, X., Tecuapetla, F., and Costa, R.M. (2014). Basal ganglia subcircuits distinctively encode the parsing and concatenation of action sequences. Nat. Neurosci. 17, 423–430.

Kato, M., and Kimura, M. (1992). Effects of reversible blockade of basal ganglia on a voluntary arm movement. J Neurophysiol 68, 1516–1534.

Kim, H.F., Amita, H., and Hikosaka, O. (2017). Indirect Pathway of Caudal Basal Ganglia for Rejection of Valueless Visual Objects. Neuron 94, 920-930.e3.

Kimura, M., Kato, M., Shimazaki, H., Watanabe, K., and Matsumoto, N. (1996). Neural information transferred from the putamen to the globus pallidus during learned movement in the monkey. J. Neurophysiol. 76, 3771–3786.

Kita, H. (2007). Globus pallidus external segment. Prog Brain Res 160, 111–133.

Kovaleski, R.F., Callahan, J.W., Chazalon, M., Wokosin, D.L., Baufreton, J., and Bevan, M.D. (2020). Dysregulation of external globus pallidus-subthalamic nucleus network dynamics in parkinsonian mice during cortical slow-wave activity and activation. J. Physiol. 598, 1897–1927.

Kravitz, A. V, and Kreitzer, A.C. (2012). Striatal mechanisms underlying movement, reinforcement, and punishment. Physiology (Bethesda). 27, 167–177.

Kravitz, A. V, Freeze, B.S., Parker, P.R., Kay, K., Thwin, M.T., Deisseroth, K., and Kreitzer, A.C. (2010). Regulation of parkinsonian motor behaviours by optogenetic control of basal ganglia circuitry. Nature 466, 622–626.

Kravitz, A. V, Tye, L.D., and Kreitzer, A.C. (2012). Distinct roles for direct and indirect pathway striatal neurons in reinforcement. Nat. Neurosci. 15, 816–818.

Kravitz, A. V, Owen, S.F., and Kreitzer, A.C. (2013). Optogenetic identification of striatal projection neuron subtypes during in vivo recordings. Brain Res 1511, 21–32.

Lalive, A.L., Lien, A.D., Roseberry, T.K., Donahue, C.H., and Kreitzer, A.C. (2018). Motor thalamus supports striatum-driven reinforcement. Elife 7, e34032.

LeBlanc, K.H., London, T.D., Szczot, I., Bocarsly, M.E., Friend, D.M., Nguyen, K.P., Mengesha, M.M., Rubinstein, M., Alvarez, V.A., and Kravitz, A. V (2020). Striatopallidal neurons control avoidance behavior in exploratory tasks. Mol. Psychiatry 25, 491–505.

Lee, J., Wang, W., and Sabatini, B.L. (2020). Anatomically segregated basal ganglia pathways allow parallel behavioral modulation. Nat. Neurosci. 23, 1388–1398.

Lenz, J.D., and Lobo, M.K. (2013). Optogenetic insights into striatal function and behavior. Behav. Brain Res. 255, 44–54.

Lilascharoen, V., Wang, E.H.-J., Do, N., Pate, S.C., Tran, A.N., Yoon, C.D., Choi, J.-H., Wang, X.-Y., Pribiag, H., Park, Y.-G., et al. (2021). Divergent pallidal pathways underlying distinct Parkinsonian behavioral deficits. Nat. Neurosci. 24, 504–515.

Lynd-Balta, E., and Haber, S.N. (1994). Primate striatonigral projections: a comparison of the sensorimotor-related striatum and the ventral striatum. J. Comp. Neurol. 345, 562–578.

Madisen, L., Zwingman, T.A., Sunkin, S.M., Oh, S.W., Zariwala, H.A., Gu, H., Ng, L.L., Palmiter, R.D., Hawrylycz, M.J., Jones, A.R., et al. (2010). A robust and high-throughput Cre reporting and characterization system for the whole mouse brain. Nat Neurosci 13, 133–140.

Mallet, N., Micklem, B.R., Henny, P., Brown, M.T., Williams, C., Bolam, J.P., Nakamura, K.C., and Magill, P.J. (2012). Dichotomous organization of the external globus pallidus. Neuron 74, 1075– 1086.

Mallet, N., Schmidt, R., Leventhal, D., Chen, F., Amer, N., Boraud, T., and Berke, J.D. (2016). Arkypallidal Cells Send a Stop Signal to Striatum. Neuron 89, 308–316.

Markowitz, J.E., Gillis, W.F., Beron, C.C., Neufeld, S.Q., Robertson, K., Bhagat, N.D., Peterson, R.E., Peterson, E., Hyun, M., Linderman, S.W., et al. (2018). The Striatum Organizes 3D Behavior via Moment-to-Moment Action Selection. Cell 174, 44-58.e17.

Mastro, K.J., Bouchard, R.S., Holt, H.A.K., and Gittis, A.H. (2014). Transgenic mouse lines subdivide external segment of the globus pallidus (GPe) neurons and reveal distinct GPe output pathways. J. Neurosci. 34.

Mastro, K.J., Zitelli, K.T., Willard, A.M., Leblanc, K.H., Kravitz, A. V., and Gittis, A.H. (2017). Cell-specific pallidal intervention induces long-lasting motor recovery in dopamine-depleted mice. Nat. Neurosci. 20, 815–823.

Matamales, M., McGovern, A.E., Mi, J.D., Mazzone, S.B., Balleine, B.W., and Bertran-Gonzalez, J. (2020). Local D2-to D1-neuron transmodulation updates goal-directed learning in the striatum. Science (80-.). 367, 549–555.

Mink, J.W. (1996). THE BASAL GANGLIA: FOCUSED SELECTION AND INHIBITION OF COMPETING MOTOR PROGRAMS. Prog. Neurobiol. 50, 381–425.

Mink, J.W., and Thach, W.T. (1991). Basal ganglia motor control.II. Late pallidal timing relative to movement onset and inconsistent pallidal coding of movement parameters. J Neurophysiol 65, 301–329.

Nambu, A., Tokuno, H., and Takada, M. (2002). Functional significance of the cortico-subthalamo-pallidal “hyperdirect” pathway. Neurosci. Res. 43, 111–117.

Nelson, A.B., and Kreitzer, A.C. (2014). Reassessing Models of Basal Ganglia Function and Dysfunction. Annu. Rev. Neurosci. 37, 117–135.

Parker, J.G., Marshall, J.D., Ahanonu, B., Wu, Y.-W., Kim, T.H., Grewe, B.F., Zhang, Y., Li, J.Z., Ding, J.B., Ehlers, M.D., et al. (2018). Diametric neural ensemble dynamics in parkinsonian and dyskinetic states. Nature 557, 177–182.

Parker, K.L., Kim, Y., Alberico, S.L., Emmons, E.B., and Narayanan, N.S. (2016). Optogenetic approaches to evaluate striatal function in animal models of Parkinson disease. Dialogues Clin. Neurosci. 18, 99–107.

Pollack, A.E., Harrison, M.B., Wooten, G.F., and Fink, J.S. (1993). Differential localization of A2a adenosine receptor mRNA with D1 and D2 dopamine receptor mRNA in striatal output pathways following a selective lesion of striatonigral neurons. Brain Res. 631, 161–166.

Polyakova, Z., Chiken, S., Hatanaka, N., and Nambu, A. (2020). Cortical control of subthalamic neuronal activity through the hyperdirect and indirect pathways in monkeys. J. Neurosci. 40, 7451–7463.

Roseberry, T.K., Lee, A.M., Lalive, A.L., Wilbrecht, L., Bonci, A., and Kreitzer, A.C. (2016). Cell-Type-Specific Control of Brainstem Locomotor Circuits by Basal Ganglia. Cell 164, 526–537.

Sales-Carbonell, C., Taouali, W., Khalki, L., Pasquet, M.O., Petit, L.F., Moreau, T., Rueda-Orozco, P.E., and Robbe, D. (2018). No Discrete Start/Stop Signals in the Dorsal Striatum of Mice Performing a Learned Action. Curr. Biol. 28, 3044-3055.e5.

Sano, H., Chiken, S., Hikida, T., Kobayashi, K., and Nambu, A. (2013). Signals through the striatopallidal indirect pathway stop movements by phasic excitation in the substantia nigra. J. Neurosci. 33, 7583–7594.

Schechtman, E., Noblejas, M.I., Mizrahi, A.D., Dauber, O., and Bergman, H. (2016). Pallidal spiking activity reflects learning dynamics and predicts performance. Proc. Natl. Acad. Sci. U. S. A. 113, E6281–E6289.

Schiffmann, S.N., Jacobs, O., and Vanderhaeghen, J.J. (1991). Striatal restricted adenosine A2 receptor (RDC8) is expressed by enkephalin but not by substance P neurons: an in situ hybridization histochemistry study. J. Neurochem. 57, 1062–1067.

Schor, J.S., and Nelson, A.B. (2019). Multiple stimulation parameters influence efficacy of deep brain stimulation in parkinsonian mice. J. Clin. Invest. 129, 3833–3838.

Smith, Y., Bevan, M.D., Shink, E., and Bolam, J.P. (1998). Microcircuitry of the direct and indirect pathways of the basal ganglia. Neuroscience 86, 353–387.

Soares-Cunha, C., Coimbra, B., Sousa, N., and Rodrigues, A.J. (2016). Reappraising striatal D1-and D2-neurons in reward and aversion. Neurosci. Biobehav. Rev. 68, 370–386.

Soares, J., Kliem, M.A., Betarbet, R., Greenamyre, J.T., Yamamoto, B., and Wichmann, T. (2004). Role of external pallidal segment in primate parkinsonism: comparison of the effects of 1-methyl-4-phenyl-1,2,3,6-tetrahydropyridine-induced parkinsonism and lesions of the external pallidal segment. J Neurosci 24, 6417–6426.

Spix, T.A., Nanivadekar, S., Toong, N., Kaplow, I.M., Isett, B.R., Goksen, Y., Pfenning, A.R., and Gittis, A.H. (2021). Population-specific neuromodulation prolongs therapeutic benefits of deep brain stimulation. Science 374, 201–206.

Taverna, S., Ilijic, E., and Surmeier, D.J. (2008). Recurrent Collateral Connections of Striatal Medium Spiny Neurons Are Disrupted in Models of Parkinson’s Disease. J. Neurosci. 28, 5504– 5512.

Tecuapetla, F., Jin, X., Lima, S.Q., and Costa, R.M. (2016). Complementary Contributions of Striatal Projection Pathways to Action Initiation and Execution. Cell 166.

Tepper, J.M., Wilson, C.J., and Koós, T. (2008). Feedforward and feedback inhibition in neostriatal GABAergic spiny neurons. Brain Res. Rev. 58, 272–281.

Turner, R.S., and Anderson, M.E. (1997). Pallidal discharge related to the kinematics of reaching movements in two dimensions. J Neurophysiol 77, 1051–1074.

Valsky, D., Heiman Grosberg, S., Israel, Z., Boraud, T., Bergman, H., and Deffains, M. (2020). What is the true discharge rate and pattern of the striatal projection neurons in Parkinson’s disease and Dystonia? Elife 9, e57445.

Wichmann, T., and DeLong, M.R. (2011). Deep-brain stimulation for basal ganglia disorders. Basal Ganglia 1, 65–77.

Yttri, E.A., and Dudman, J.T. (2016). Opponent and bidirectional control of movement velocity in the basal ganglia. Nature 533, 402–406.

Yu, C., Cassar, I.R., Sambangi, J., and Grill, W.M. (2020). Frequency-Specific Optogenetic Deep Brain Stimulation of Subthalamic Nucleus Improves Parkinsonian Motor Behaviors. J. Neurosci. 40, 4323–4334.

